# L-DOS47 enhances response to immunotherapy in pancreatic cancer tumor

**DOI:** 10.1101/2023.08.28.555194

**Authors:** Bruna Victorasso Jardim-Perassi, Pietro Irrera, Dominique Abrahams, Veronica C. Estrella, Bryce Ordway, Samantha R. Byrne, Andrew A. Ojeda, Christopher J. Whelan, Jongphil Kim, Matthew S. Beatty, Sultan Damgaci-Erturk, Dario Livio Longo, Kim J. Gaspar, Gabrielle M. Siegers, Barbara A. Centeno, Justin Y.C. Lau, Arig Ibrahim-Hashim, Shari A. Pilon-Thomas, Robert J. Gillies

**Affiliations:** Department of Metabolism and Physiology, H. Lee Moffitt Cancer Center and Research Institute, Tampa, FL, USA; University of South Florida, Comparative Medicine, Tampa, FL, USA; Massachusetts General Hospital, Harvard Medical School, Boston, Massachusetts; Department of Immunology, H. Lee Moffitt Cancer Center and Research Institute, Tampa, FL, USA; Department of Biological Sciences, University of Illinois, Chicago, IL, USA; Department of Biostatistics and Bioinformatics, H. Lee Moffitt Cancer Center and Research Institute, Tampa, FL, USA; Merck &Co, 770 Sumneytown Pike, West Point, PA, USA; Institute of Biostructures and Bioimaging (IBB), National Research Council of Italy (CNR), Turin, Italy; Helix BioPharma Corp., Toronto, ON, Canada; Department of Pathology, H. Lee Moffitt Cancer Center and Research Institute, Tampa, FL, USA; Small Animal Imaging Laboratory (SAIL), H. Lee Moffitt Cancer Center and Research Institute, Tampa, FL, USA; GE HealthCare, Niskayuna, NY, USA

**Keywords:** pancreatic cancer, acidosis, immunotherapy, L-DOS47

## Abstract

Acidosis is an important immunosuppressive mechanism that leads to tumor growth. Therefore, we investigated the neutralization of tumor acidity to improve immunotherapy response. L-DOS47, a new targeted urease immunoconjugate designed to neutralize tumor acidity, has been well tolerated in phase I/IIa trials. L-DOS47 binds CEACAM6, a cell surface protein highly expressed in gastrointestinal cancers, allowing urease to cleave endogenous urea into two NH4+ and one CO2, thereby raising local pH. To test the synergetic effect of neutralizing tumor acidity with immunotherapy, we developed a pancreatic orthotopic murine tumor model (KPC961) expressing human CEACAM6. Our results demonstrate that combining L DOS47 with anti-PD1 significantly increases the efficacy of anti-PD1 monotherapy, reducing tumor growth for up to 4 weeks.

## Introduction

Acidosis is a well-established feature of cancer, favoring its progression and metastatic spread. Tumor acidosis is an extracellular condition provoked by the increased production of acidic molecules and protons, it alters the neoplastic tissue’s physiological homeostasis, impacting many cell subtypes, including macrophages, T-and-B-cells, epithelial cells, and fibroblasts [1, 2]. In healthy conditions, acidosis is also exploited, particularly in lymph nodes where T cells sustain an acidic environment to suppress their own functions and become active only after they leave the acidic lymph node site [3]. As such, immune suppression and deregulation represent one of the crucial processes to be addressed when treating cancer because these events occur as physiological responses to the acidosis condition [4-6].

An acidic environment can contribute to immune impairment, where many components, such as macrophages [7], T cells [8], and natural killer (NK) cells shift to a state that supports tumor growth [9]. Therapies inhibiting these pathways can restore immune responses against cancer; however, if the underlying acidic condition remains, these effects are lost even when combined with immunotherapies [7].

Immune treatments and inhibitory strategies alone are often insufficient to overcome the effects of acidosis. Directly counteracting acidity through alkalinization treatments is a more reliable approach and coupling buffer therapies with immune/inhibitory treatments may result in better outcomes [10-12]. Studies have shown how this strategy could be applied in several tumor models: combining acidity-lowering drugs with chemotherapy was essential to overcome chemoresistance in human and murine tumors [13-16], and chronic oral administration of bicarbonate coupled with immune therapy significantly reduced tumor growth compared to the monotherapies alone [17]. Yet, even if these approaches are proven successful in the preclinical setting, a direct translation into the clinic is often delayed or unfeasible because they are either not effective in patients [18] or not safe enough for long term use.

In the present study, we used a combination approach involving L-DOS47, a new pH-targeting molecule developed by Helix BioPharma (Toronto, ON, Canada), in addition to the canonical immune therapy provided by the administration of an anti-programmed death (PD1) antibody. L-DOS47 is an immunoconjugate comprising multiple copies of a camelid single domain antibody that specifically binds the carcinoembryonic antigen-related cell adhesion molecule 6 (CEACAM6) constitutively upregulated in human cancer cells [19, 20], conjugated to a jack bean derived urease enzyme [21]. Its mechanism of action involves selective binding to CEACAM6 on the tumor cell surface, localizing the urease, which converts endogenous urea into NH_3_ and CO_2_ with the net production of bicarbonate and hydroxyl ions and causes alkalinization of the extracellular tumor microenvironment (TME). In Phase I clinical trials, L-DOS47 was well tolerated in patients with non-small-cell lung cancer (NSCLC) and yielded encouraging results when combined with pemetrexed plus carboplatin, with 75% of patients showing clinical benefit (stable disease, complete or partial response) [22]. Since L-DOS47 is suitable for cancers that express high levels of its target antigen CEACAM6, pancreatic and gastrointestinal cancers are potential other applications in addition to lung cancer [23, 24].

## Results

### Development of an orthotopic pancreatic tumor model that expresses human CEACAM6

An orthotopic pancreatic tumor model was generated using the murine PDAC cell line KPC961, which was engineered to express human CEACAM6 (hCEACAM6). After transduction, flow cytometric analyses confirmed the presence of 97.7% hCEACAM6-expressing cells in clone 1B6 **(Fig. 1A; Fig S1)**. Metabolic profiling showed no differences in energy metabolism between the KPC961 parental and clone 1B6 cells (**Fig 1B-C**).

**Figure 1.**
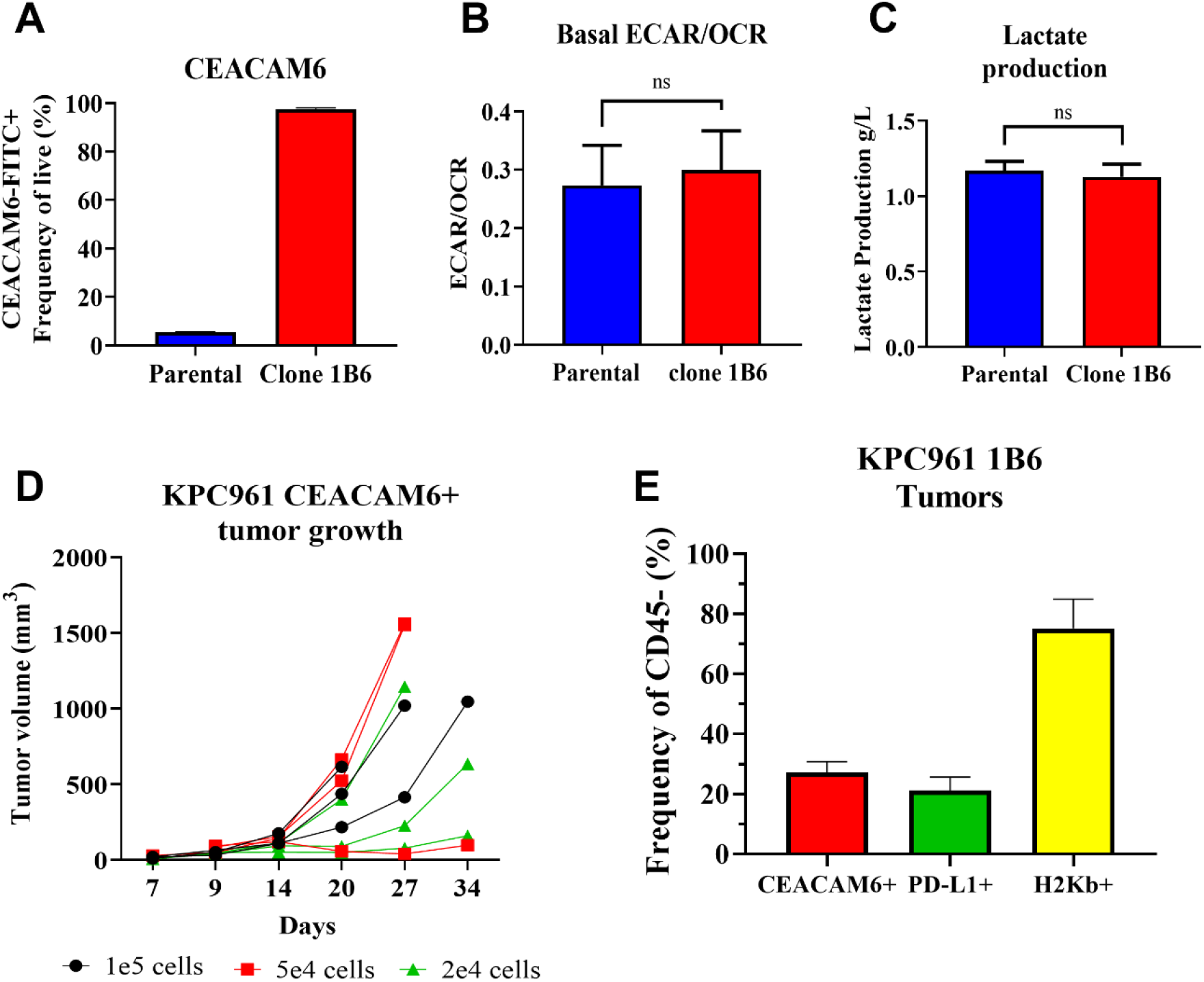
Generation of an orthotopic pancreatic adenocarcinoma murine tumor model expressing human CEACAM6. **A**. Percentage of CEACAM6-expressing cells analyzed by flow cytometry in KPC961 parental and KPC961 clone 1B6 cells. **B**. Ratio of extracellular acidification (ECAR) and oxygen consumption rate (OCR) profiles in KPC961 parental and clone 1B6 cells; **C**. Lactate production in KPC961 parental and clone 1B6 cells. **D**. Tumor growth as a function of inoculum size. KPC961-CEACAM6 cells were inoculated into the pancreas of B6.129 mice at 100000, 50000, and 20000 cells, and tumor volumes were measured by ultrasound. Ascites fluid was observed in two mice inoculated with 100,000 cells on days 24 and 29, two mice inoculated with 50,000 cells at day 29 and one mouse inoculated with 20,000 cells at day 27. **E**. Expression of CEACAM6, PD-L1 and H2Kb in KPC961-1B6 cells of inoculated orthotopic tumors by flow cytometry.

**Figure 2.**
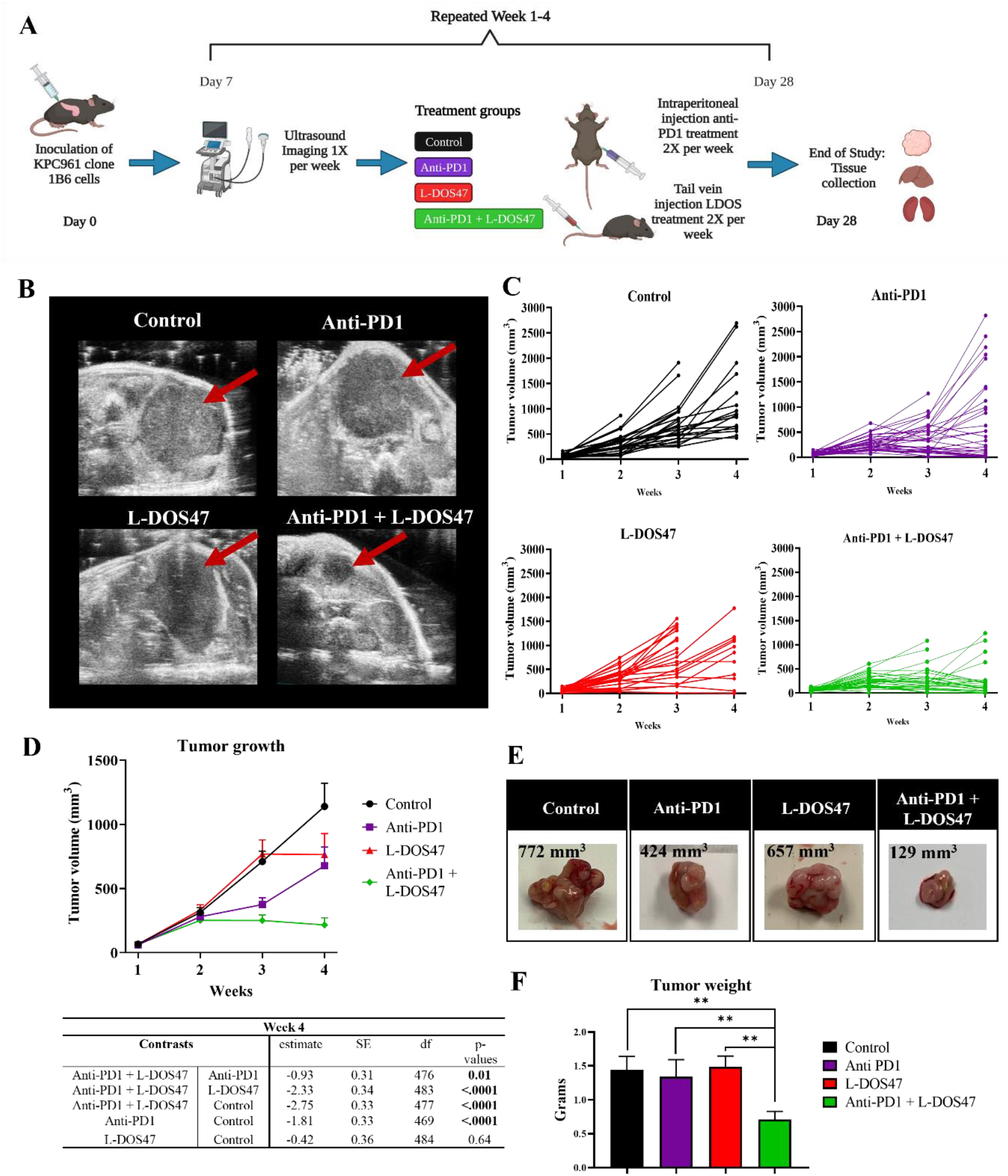
Therapeutic efficacy in KPC961-1B6 orthotopic tumors. **A**. Experimental design. **B**. Representative ultrasound images for KPC961-1B6 pancreatic tumors. **C**. Individual tumor growth for each therapy group for the 3 replicates (n=28 in control group; n=35 in anti-PD1 group; n=23 in L-DOS47 group; n=35 in anti-PD1 + L-DOS47). **D**. Mean tumor volume ± SEM for each therapy group with corresponding table showing the differences at end point (week 4). **E**. Representative *ex vivo* images of KPC961 1B6 tumors for each therapy group with corresponding tumor volumes as measured with US (rounded to integral digit). **F**. Mean tumor weight at endpoint. Statistical significance is reported based on *p-values* obtained from the Tukey’s test (see Table 2).

We additionally confirmed that the KPC961 clone 1B6 could form orthotopic tumors in immunocompetent B6.129 mice **(Fig. 1D)** and showed continued CEACAM6 expression (27.2 ± 3.56 %) in inoculated tumors as indicated by flow cytometry **(Fig. 1E; Fig S2)**. These tumors were also tested for the presence of the major histocompatibility complex (MHC) class I molecule H2K^b^ and PD-1 ligand 1 (PD-L1), representing 75.2 ± 9.8 % and 21.33 ± 4.45 % of the tumor cells respectively (**Fig 1E; Fig S2**).

### L-DOS47 has a synergistic effect on anti-PD1 therapy in reducing tumor growth

Once pharmacodynamic studies confirmed that L-DOS47 increases tumor pHe in the KPC961-1B6 orthotopic model, we investigated the therapeutic efficacy of L-DOS47 as a monotherapy and in combination with anti-PD1. Immunocompetent mice were inoculated orthotopically, and after tumor establishment (week 1, day 6-7), mice were randomized into groups with equal tumor volume averages before initiating therapies. Mice were treated twice a week, and tumor volumes were measured weekly by ultrasound (US) imaging to monitor tumor growth (**Fig. 4A**).

Linear mixed-effects models were used to test for differences in tumor growth among treatment arms. Tumor growth for all treatment arms was significantly greater than for the combination treatment of anti-PD1 + L-DOS47 (**Table S1**). Statistical analyses showed that the combination of anti-PD1 + L-DOS47 significantly reduced tumor growth when compared with control (p=0.01) and L-DOS47 monotherapy (p=0.04) groups at week 2 (days 13-14), and it differed significantly from anti-PD1 monotherapy at weeks 3 and 4 (days 20-21 and 27-28). Although anti-PD1 monotherapy was effective in reducing tumor growth after 3 weeks when compared with the control group (p<0.0001), the combination of anti-PD1 + L-DOS47 showed the greatest efficacy and had a synergistic effect when compared with the anti-PD1 monotherapy (p=0.03 for week 2 and p=0.01 for week 3) (**Figure 4B-D; Table 1**). Tumor growth plots for each experimental replicate are shown in **Fig. S4**. In addition, tumor weights measured at endpoint (week 4) were significantly lower in the anti-PD1 + L-DOS47 group when comparing with control (p=0.01), L-DOS47 (p=0.01) and anti-PD1 (p=0.03) monotherapies (**Fig 4E-F; Table 2**).

**Table 1.** Post-hoc comparison of estimated marginal means based on the linear mixed model for the tumor growth of each therapy group (combined replicates 1, 2 and 3).

| Day 13 |  |  |  |  |  |
| --- | --- | --- | --- | --- | --- |
| Contrasts |  | estimate | SE | df | p-values |
| Anti-PD1 + L-DOS47 | Anti-PD1 | -0.26 | 0.22 | 353 | 0.65 |
| Anti-PD1 + L-DOS47 | L-DOS47 | -0.66 | 0.25 | 405 | **0.04** |
| Anti-PD1 + L-DOS47 | Control | -0.74 | 0.24 | 363 | **0.01** |
| Anti-PD1 | Control | -0.48 | 0.25 | 334 | 0.21 |
| L-DOS47 | Control | -0.08 | 0.26 | 432 | 0.98 |
| Day 21 |  |  |  |  |  |
| Contrasts |  | estimate | SE | df | p-values |
| Anti-PD1 + L-DOS47 | Anti-PD1 | -0.60 | 0.22 | 340 | **0.03** |
| Anti-PD1 + L-DOS47 | L-DOS47 | -1.49 | 0.24 | 389 | **<.0001** |
| Anti-PD1 + L-DOS47 | Control | -1.75 | 0.24 | 347 | **<.0001** |
| Anti-PD1 | Control | -1.15 | 0.24 | 316 | **<.0001** |
| L-DOS47 | Control | -0.25 | 0.25 | 418 | 0.75 |
| Day 27 |  |  |  |  |  |
| Contrasts |  | estimate | SE | df | p-values |
| Anti-PD1 + L-DOS47 | Anti-PD1 | -0.93 | 0.31 | 476 | **0.01** |
| Anti-PD1 + L-DOS47 | L-DOS47 | -2.33 | 0.34 | 483 | **<.0001** |
| Anti-PD1 + L-DOS47 | Control | -2.75 | 0.33 | 477 | **<.0001** |
| Anti-PD1 | Control | -1.81 | 0.33 | 469 | **<.0001** |
| L-DOS47 | Control | -0.42 | 0.36 | 484 | 0.64 |

Degrees-of-freedom method: kenward-roger
Results are given on the log2 (not the response) scale.
P value adjustment: tukey method for comparing a family of 4 estimates

**Table 2.**
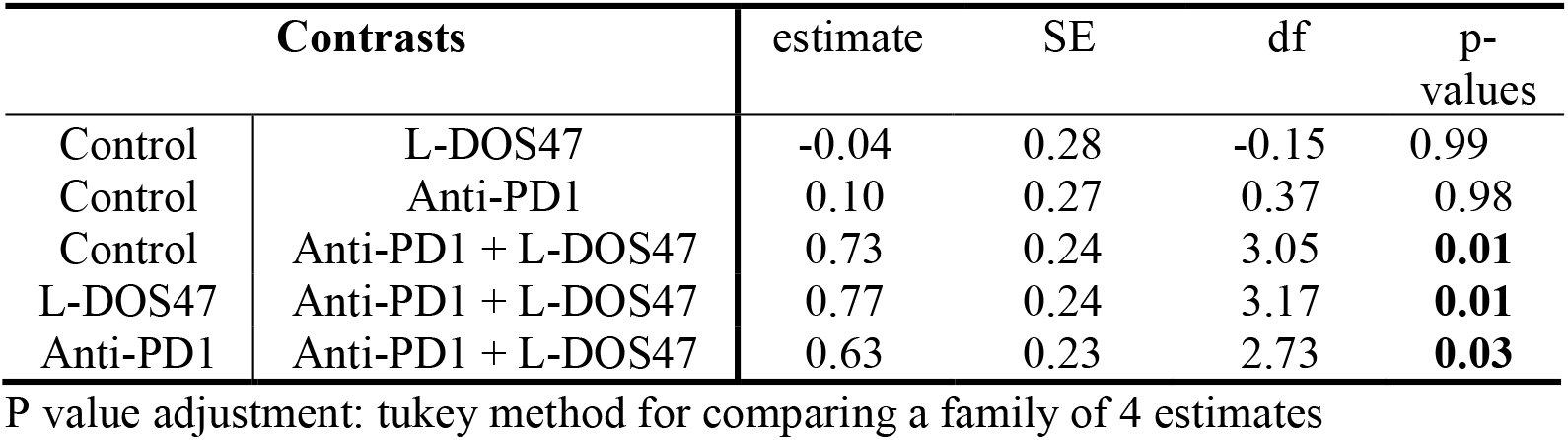
Tukey’s Honest Significant Difference post-hoc tests, and pairwise comparisons of tumor weight marginal means.

| Contrasts |  | estimate | SE | df | p-values |
| --- | --- | --- | --- | --- | --- |
| Control | L-DOS47 | -0.04 | 0.28 | -0.15 | 0.99 |
| Control | Anti-PD1 | 0.10 | 0.27 | 0.37 | 0.98 |
| Control | Anti-PD1 + L-DOS47 | 0.73 | 0.24 | 3.05 | **0.01** |
| L-DOS47 | Anti-PD1 + L-DOS47 | 0.77 | 0.24 | 3.17 | **0.01** |
| Anti-PD1 | Anti-PD1 + L-DOS47 | 0.63 | 0.23 | 2.73 | **0.03** |

P value adjustment: tukey method for comparing a family of 4 estimates

## Discussion

Although major advances have been made in other solid tumors, the utility of immunotherapy in PDAC has yet to be demonstrated. Various immunotherapies that have been tried have proven largely unsuccessful, likely due to PDAC’s characteristically low tumor mutational burden and highly immunosuppressive tumor microenvironment [25].

Pembrolizumab, an anti-PD1 immune checkpoint inhibitor, has been approved by the FDA for a subset of patients with advanced PDAC whose tumors have been identified as mismatch repair deficient (dMMR) or microsatellite instability high (MSI-H), the latter of which accounts for only 0.8 to 2% of patients [25]. In the context of KEYNOTE-158, a recent study of pembrolizumab in “all comers” in solid tumors, an 18.2% overall response rate (ORR), 4.0 months median overall survival (OS) and 13.4 months median duration of response were reported with only three partial responses and one complete response in 22 PDAC patients with MSI-H or dMMR deficiency [26]. In a first dual approach with 65 patients with recurrent or metastatic PDAC, the PD-L1 antibody durvalumab in combination with the anti-cytotoxic T-lymphocyte antigen-4 (CTLA-4) antibody tremelimumab yielded only a 3.1% overall response rate compared to 0% for durvalumab monotherapy [27].

PDAC has been historically considered immunologically “cold”, yet a growing number of studies indicate inherent heterogeneity and potential reversibility of this phenotype [28]. Attempting to improve outcomes in PDAC by utilizing a combination of checkpoint inhibitors with therapies targeting immunosuppressive features, such as acidosis in the tumor microenvironment, is a logical next step.

Acidosis is one of the major drivers and supporters of immune impairment during neoplastic formation and it is imperative to target this for a successful treatment outcome [29]. Here we show in a murine orthotopic PDAC model that L-DOS47, an immunoconjugate that binds specifically to CEACAM6-expressing tumor cells, counters acidosis by raising pHe locally through the ureolytic activity of its urease enzyme moiety. L-DOS47 also acts synergistically with anti-PD1 therapy to slow tumor growth in this model.

Previously, the specificity and cytotoxicity of L-DOS47 were confirmed in different CEACAM6-expressing cancer cell lines (BxPC-3 pancreatic, A549 lung, MCF7 breast, and CEACAM6-transfected H23 lung), where the response to L-DOS47 was positively correlated with levels of CEACAM6 expression [21]. L-DOS47 was most effective in reducing *in vitro* viability of BxPC3 cells, which had the greatest levels of CEACAM6 expression [21]. Furthermore, tumor growth was significantly inhibited by L-DOS47 in a xenograft tumor model using BxPC3 [21]. This xenograft model utilized immunocompromised nude mice, and so any impact of L-DOS47 on the immune system could not be assessed [21]. The model we have now developed using hCEACAM6-expressing KPC961 cells has overcome this limitation, and enabled observation of the profound enhancement of anti-PD1 efficacy when combined with L-DOS47.

Cellular metabolism was not altered during model establishment, and no spontaneous tumor rejection was observed in control arms of our *in vivo* experiments, showing that no undue immunogenicity was caused by expression of the human CEACAM6 protein.

L-DOS47 was designed as a novel variation on antibody directed enzyme prodrug therapy (ADEPT), in which the antibody component targets the molecule to its antigen – in this case CEACAM6 -on the tumor cell surface; however, unlike conventional ADEPT, in which prodrugs are administered systemically for the enzyme component to act upon, L-DOS47 uses the metabolite urea as a substrate, which is constitutively present in tumor tissues [30]. The urease enzyme component of L-DOS47 converts endogenous extracellular urea into ammonia and CO_2_ resulting in the formation of bicarbonate and hydroxyl ions, which alkalize the tumor microenvironment [21].

In therapeutic efficacy studies *in vivo*, optimal effects were observed with twice-weekly administration of L-DOS47 with anti-PD1, as the combination group exhibited significantly lower tumor volumes and weights compared to either monotherapy group. A significant difference was found at week 2 when comparing the combination therapy to the control and the L-DOS47 monotherapy groups. Strikingly, L-DOS47 together with anti-PD-1 was significantly more effective in controlling tumor growth than anti-PD1 alone. In the experiments shown here, anti-PD1 was administered 4 hours before L-DOS47; however, in a separate *in vivo* experiment, we found no significant differences in outcome comparing whether L-DOS47 or anti-PD1 was administered first (data not shown). Since anti-PD1 is known to have a long circulating half-life, and dosing was done twice weekly, anti-PD1 would have been present in the mice as of the first treatment and, as such, if a difference was to be found, it would only be applicable to the first dose.

Surprisingly, anti-PD1 monotherapy also provided some efficacy in our *in vivo* model, which is not commonly observed in patients with pancreatic cancer [31]. However, some patients with MSI-H tumors eventually respond to anti-PD1 with similar survival to that achieved with standard treatment for pancreatic cancer [31]. In line with this, it has been postulated that the immunologic *status quo* of the pancreatic tumor microenvironment can drastically differ across patients, where a pro-inflammatory state seems to adjuvate anti-PD1 efficacy [32, 33]. While additional studies are warranted to evaluate the effects of L-DOS47 administration on immune cell subsets in the tumor microenvironment, it is nonetheless clear that the combination of L-DOS47 with anti-PD1 remained the most effective treatment overall in this study.

Other preclinical studies have demonstrated that buffer systems such as sodium bicarbonate can alkalinize tumor pH and reverse the consequences of acidosis [18, 34]. Oral administration of sodium bicarbonate prevented tumor development [35] and reduced invasion and metastasis in various tumor models, although it had no effect on primary tumor growth [11, 36-38]. We have also previously shown that buffering tumor pH using sodium bicarbonate can improve the antitumor response to immune checkpoint therapies, as well as the adoptive transfer of T lymphocytes in B16 melanoma and Panc02 pancreatic tumor models. Combination therapy with bicarbonate and anti-PD1 or anti-CTLA-4 impaired tumor growth and led, in some cases, to tumor regression [17] .

Unfortunately, these preclinical findings have not been supported by positive clinical trial results. The first three clinical trials using oral sodium bicarbonate (NCT01350583, NCT01198821, NCT01846429) failed mainly due to poor patient compliance associated with its unpleasant taste and/or gastrointestinal side effects [18, 39]. More recently, patients with advanced pancreatic cancer (UMIN000035659) [40] and small cell lung cancer (UMIN000043056) [41] did show improved outcomes when receiving alkalization treatment, which included an alkaline diet and/or oral sodium bicarbonate (3.0-5.0 g/day). However, these were retrospective analyses of non-randomized single-center studies that included only a small number of patients [42].

Conversely, L-DOS47 has been proven safe and well tolerated in Phase I/IIa clinical trials in heavily pretreated NSCLC patients both as a monotherapy and in combination with pemetrexed and carboplatin [22, 43]. Encouraging progression-free survival and clinical benefit were observed, particularly in patients also receiving pemetrexed/carboplatin (41.7% objective response rate and 75% clinical benefit). Additionally, a number of patients continued on L-DOS47 monotherapy well past the four protocol-mandated cycles. The maximum tolerated dose was not reached in either completed study at doses up to 13.55 μg/kg in the monotherapy and 9.0 μg/kg in the combination therapy study, which are both above the human equivalent dose of L-DOS47 used in the current study (7.3 μg/kg).

With high expression of CEACAM6, L-DOS47 could be an ideal solution for treating cancers such as lung, gastric, colorectal and pancreatic in combination with chemo-, immuno-, and radio-therapies, as well as other therapeutic modalities including cell and oncolytic virus therapies where acidosis is a limiting factor for efficacy [23, 24]. Indeed, a Phase I/II trial is currently ongoing to evaluate L-DOS47 in combination with doxorubicin in advanced pancreatic cancer patients (NCT04203641).

In this preclinical study, the reduction of tumor acidosis with L-DOS47 strengthened the anti-tumor response to anti-PD1 treatment by providing significantly greater tumor control. Future studies will test the efficacy of combining L-DOS47 and anti-PD1 or other immunotherapies in additional CEACAM6-expressing preclinical cancer models and clinical trials. In conclusion, L-DOS47 offers significant potential for broad applicability in combination with a growing number of innovative cancer treatments in the future.

## Materials and Methods

### 1. Transduction and selection of CEACAM6 expressing clone

The murine pancreatic cancer cell line UN-KPC-961 (KPC961) was obtained via MTA from Dr. Surinder K. Batra (University of Nebraska Medical Center, Omaha, NE) [44]. Cells were retrovirally infected with hCEACAM6 pLenti-GIII-EF1a lentivirus and a CEACAM6 expressing subclone was selected for this study. For the transduction, cells were trypsinized, centrifuged, and diluted to a concentration of 50K cells per mL in DMEM/F12 media (Gibco, Waltham, MA) supplemented with 10% FBS, 1% of P/S (Sigma, St. Louis, MO) and containing 5μg/ml polybrene. 8 μL of the CEACAM6 lentivirus was added to 500 μL of complete DMEM/F12, and the lentiviral mixture was added to the wells of a 6-well plate. 1 mL of the previously diluted cells was added on top of the lentiviral mixture in the 6-well plate. 18 hours after addition of the cells, the media was removed, and cells were cultured in complete DMEM/F12. After 24 hours, the media was replaced with complete DMEM/F12 containing 5 μg/mL puromycin (Gibco, Waltham, MA). Once at 90% confluence in the 6-well plate, cells were trypsinized and transferred to a T-75 flask to be maintained in an incubator at 37°C and 5% CO_2_.

For the selection process, cells were trypsinized, centrifuged, diluted, counted, and then passed through a 4 μm filter. Cells were diluted and counted again, and then diluted once more to achieve a final concentration of 500 cells per mL. 2 μL of the diluted cells were added to each well of 24-well plates. Drops were assessed for the presence of cells, with wells containing more than one cell omitted. All wells were then filled with DMEM/F12 containing 5μg/mL puromycin. After 24 hours and again after 120 hours, wells were visually assessed to determine which wells contained single colonies of cells. Wells containing no cells or multiple colonies were omitted. Each clone (single cell origin) was transferred to 6-well plates to be expanded. Once confluent, the cells were collected and the expression of human CEACAM6 was verified by flow cytometry.

### 2. Flow cytometry

#### a. CEACAM6 expression

KPC961-CEACAM6-transduced subclones and KPC961 parental cells (negative control) were resuspended in FACS buffer (PBS with 5% FBS, 1 mM EDTA and 0.1% Sodium Azide) to a concentration of 0.5− 1×10^6^ cells/mL for flow cytometric analysis. Cells were stained in FACS buffer with anti-CEACAM6-FITC antibody (Sino Biological 10823-R408R, Sino Biological, Inc., Wayne, PA) at 10 μg/mL for 30 min at 4°C in the dark. After incubation, cells were washed with FACS buffer, centrifuged, and resuspended in FACS Buffer containing 1.25 μg/mL live/dead PI reagent (Bioscience Propidium Iodide - Fisher 5018262, Fisher Scientific, Waltham, MA). Flow cytometry data were acquired using the BD FACSCelesta (BD Biosciences, Franklin Lakes, NJ) and analyzed with FlowJo ver10.8.1 (Tree Star, Ashland, OR).

#### b. Tumor Digestion and Single cell preparation for Flow Cytometry Analysis

Single cell suspensions were prepared from KPC961-1B6 tumors by first cutting the tumor into minute fragments. These fragments were placed in a GentleMACS C-tube and processed by enzymatic digestion in HBSS (Life Technologies, Carlsbad, CA) containing 1 mg/mL Collagenase D, 1 mg/mL DNAse I and 2.5 mg/mL Hyaluronidase (all from Sigma-Aldrich) and dissociated using the GentleMACS Dissociator (Miltenyi Biotec, Bergish Gladback, Germany). Following dissociation, the C-tube was stirred in a water bath at 37°C for 45 minutes. After stirring, the tissues were passed through the GentleMACS dissociator once more. The resulting suspension was put through a 70 μm cell strainer. Cells were then pelleted by centrifugation; the supernatant was discarded, and a red blood cell lysis buffer (BioLegend, San Diego, CA) was added to remove any RBCs. After RBC lysis, the cells were passed through a 70 μm cell strainer once more. Cells were then washed with PBS, pelleted, and resuspended in FACS buffer (PBS with 5% FBS, 1 mM EDTA and 0.1% Sodium Azide) at a concentration of 0.5-1 × 10^6^ cells/mL for flow cytometry analysis. Cells were labeled with the following antibodies:(CEACAM6-FITC at 10 μg/mL, Sino Biological 10823-R408R; H2Kb-Pac Blue at 0.5 mg/mL, BioLegend 116514; CD45-BV605 at 0.2 mg/mL, BioLegend 103155; and PD-L1 – PE at 0.2 mg/mL, Invitrogen 12-5982-82) in FACS buffer for 20 minutes at 4°C in the dark. Prior to antibody staining, a Fc receptor blocker (Tonbo Biosciences 70-0161-M001, Tonbo Biosciences, San Diego, CA) was added for 10 minutes at 4°C to prevent non-specific binding of antibodies. Live/dead fixable near-IR reactive dyes (Thermo Fisher Scientific, Waltham, MA) were used to exclude dead cells before analysis. Cell data were acquired using the BD FACS Celesta (BD Biosciences) and analyzed using FlowJo (Tree Star).

### 3. *In vitro* metabolic profiling

#### a. Oxygen consumption and extracellular acidification measurements

The Seahorse Extracellular Flux (XF-96) analyzer (Seahorse Bioscience, Chicopee, MA) was used to measure real-time basal oxygen consumption (OCR) and extracellular acidification rates (ECAR) in KPC961-1B6 and KPC961 parental cells. The cells were seeded in an XFe-96 microplate (Seahorse, V3-PET, 101,104–004) in normal growth media overnight. The growth media was replaced with DMEM powder base media (Sigma D5030) supplemented with 1.85 g/L sodium chloride, 1mM glutamine, and pH was set to 7.4. When testing glycolysis, cells were incubated in glucose free media and incubated for 1 hour in a non-CO_2_ incubator prior to measurement. ECAR and OCR were measured in the absence of glucose associated with the non-glycolytic activity, followed by two sequential injections of D-glucose (6 mM) and oligomycin (1 μM) in real time, which are associated with glycolytic activity (glucose-induced ECAR) and glycolytic capacity (reserve). The mitochondrial stress test was also used where cells were incubated in glucose (5.5 mM) and glutamine (1 mM) containing media and basal OCR and ECAR measured, prior to sequential injection of oligomycin (1 μM), associated with ATP-linked OCR, FCCP (1 μM) associated with mitochondrial reserve capacity and Rotenone/Antimycin A (1 μM).

Once the Seahorse assay was finalized, cells were stained using a 1:1000 dilution of the HCS NuclearMask Red Stain (Molecular Probes, cat# H10326, Molecular Probes, Eugene, OR). Cells were incubated at 37°C for 30 minutes, then washed and later imaged using the Incucyte S3 Live-Cell Analysis System (Sartorius, Gottingen, Germany). Plates were scanned using the 96-well TPP plate setting, magnification was set at 4X, and both the phase and Red FL filters were applied. The Incucyte Basic Analyzer module (version 2022B) with Top Hat background subtraction and intensity/size thresholding was used to identify the Red FL nuclei and determine both cell count and confluency. The OCR and ECAR values were normalized to cell number using cell quantification software from Incucyte S3 mentioned above.

#### b. Lactate measurement

Twenty thousand cells were seeded in a 96-well plate in 200 μL growth media containing 10% FBS. The media was collected following 48 h incubation period and measured for lactate production using the biochemistry analyzer, YSI 2900 (Xylem, Washington, D.C.). The cell densities per well were determined by Incucyte cell count application with the use of a nuclear staining technique. Cells were incubated for 30 minutes with a 1:1000 dilution of the HCS Nuclear Mask Red stain (Molecular Probes cat# H10326). After washing the cells in 1X PBS, cells were transferred to the Incucyte S3 Live-Cell Analysis System (Sartorius) where cell count and confluency was determined. Data were normalized to cell number and were reported as lactate production in g/L/ cells.

### 4. KPC961 orthotopic tumor model

Animal experiments were approved by the Institutional Animal Care and Use Committee (IACUC,protocols #8596 and #10942). Mice were obtained from The Jackson Laboratory (Bar Harbor, ME) and housed in a facility under pathogen-free conditions in accordance with IACUC standards of care at the H. Lee Moffitt Cancer Center.

KPC961-1B6 cells were inoculated orthotopically into the pancreas of B6.129 mice. Mice were dosed with Meloxicam (5 mg/kg) 30 minutes before surgery to provide analgesia, and isoflurane (2% given in 1.5 L/min oxygen breathing) was utilized to induce anesthesia during the procedure. After removing hair and sterilizing the mice midsection, abdominal skin and muscle were incised to allow a direct injection of 20 μL bolus of cells/PBS into the head of the pancreas. Closure of the abdominal cavity was accomplished in two layers and the skin layer was closed in a simple interrupted pattern with non-absorbable surgical staples, which were removed 10 days post-inoculation. The surgical procedure is described in detail in [45].

### 5. Treatments for efficacy study

Three biological replicates were performed for efficacy studies. For each experiment, B6.129 mice were inoculated orthotopically into the pancreas with KPC961-1B6 cells. Six days after tumor inoculation, tumor volumes were measured by ultrasound imaging and mice were randomized into four groups with equal tumor volume averages to initiate therapies. Mice that developed an additional tumor in the abdominal wall were excluded from the study. Treatment groups included: 1) Control (no therapy); 2) monotherapy with anti-PD1; 3) monotherapy with L-DOS47, and 4) anti-PD1 + L-DOS47 combination therapy. Treatments were administrated twice a week. Anti-PD1 (InVivoMab anti-mouse PD-1 (CD279), Clone: RMP1-14, Isotype: rat IgG2a, Bio X Cell, Lebanon, NH) was administered IP at the dose of 300 μg, while L-DOS47 was administered intravenously at 90 μg/kg. Mice in the anti-PD1 + L-DOS47 combination group were treated with anti-PD1 (300 μg) first and L-DOS47 (90 μg/kg) 4h later. Mice used in biological replicates were as follows: 39 mice in replicate 1 [Control (n=10), anti-PD-1 (n=10), L-DOS47 (n=10), anti-PD1 + L-DOS47 (n=9)]; 39 mice in replicate 2 [Control (n=9), anti-PD1 (n=10), L-DOS47 (n=8), anti-PD1 + L-DOS47 (n=12)]; and 41 mice in replicate 3 [Control (n=7), anti-PD1 (n=15), L-DOS47 (n=5), anti-PD1 + L-DOS47 (n=14)].

Tumor volumes were measured weekly by ultrasound imaging and tumor growth was monitored for 4 weeks. Mice with tumors that reached or exceeded 750 mm^3^ (end-point tumor volume) were humanely euthanized.

### 6. Ultrasound imaging

Mice were imaged with the Vevo 2100 ultrasound system (FUJIFILM VisualSonics Inc., Toronto, Canada) to measure volumes of the orthotopic pancreatic tumors. Mice were anesthetized with 1.5%-3% isoflurane delivered via nose-cone manifold, depilated with Nair, and positioned with surgical tape onto a thermo-regulated stage where electrodes and rectal probe continuously monitored their body temperature, heart rate, and respiration rate. An adjustable heat lamp and pre-warmed ultrasound gel were used to ensure that animals maintained body temperature during scanning. Scans were conducted at thicknesses of 0.05 mm with the 3D motor attachment. Regions of interest (ROIs) were obtained at parallel slices to measure tumor volume using the Vevo LAB 5.5.0 software.

### 7. Statistical analyses

Linear mixed effects models were used to test for differences in tumor growth rates among treatment arms up to 4 weeks. Tumor volume measurements were obtained at day 6-7 (week 1), 13-14 (week 2), 20-21 (week 3) and 27-28 (week 4). A tumor volume of 750 mm^3^ was considered the “endpoint”, thus if a mouse reached the endpoint tumor volume before the end of the experiment, from that time until the experiment was concluded, the tumor volume used for analyses was recorded at 750 mm^3^. Statistical analysis was carried out with package *LME4* [46] and package *lmerTEST* [47] in the R programming language [48]. Post-hoc pairwise tests of estimated marginal means based on the linear mixed model were conducted with the R *emmeans* package [49] [https://CRAN.R-project.org/package=emmeans, R. V. Lenth 2022]. Data were transformed with log base 2 to linearize the growth rate trends. Data from replicates 1, 2 and 3 were combined and we modeled log_2_ (tumor growth) as the dependent variable, day and treatment arm as the fixed main effects, the interaction of day and treatment arm, and mouse as the random effect (random intercept).

## Acknowledgment

This work has been supported in part by the Biostatistics Core, the Flow Cytometry Core Facility, and the SAIL Core Facility at the H. Lee Moffitt Cancer Center & Research Institute and NCI designated Comprehensive Cancer Center (P30-CA076292). The authors would like to thank Atul Deshpande, Frank G. Renshaw, and Christof Böhler from Helix-BioPharma for their scientific assistance. L-DOS47 was provided by Helix-Biopharma under Sponsored Research Agreements with Moffitt Cancer Center. The research reported herein was supported by the NIH/NCI (R01CA239219) and by the Associazione Italiana Ricerca Cancro (AIRC MFAG 2017 -ID 20153 project - to Dario Livio Longo).

## Data availability

All materials (data and images) reported in this article are available within the paper and its supplementary materials.

## References

1. Gatenby, R.A. and R.J. Gillies, Why do cancers have high aerobic glycolysis? Nat Rev Cancer, 2004. 4(11): p. 891–9.

2. Corbet, C. and O. Feron, Tumour acidosis: from the passenger to the driver’s seat. Nat Rev Cancer, 2017. 17(10): p. 577–593.

3. Wu, H., et al., T-cells produce acidic niches in lymph nodes to suppress their own effector functions. Nat Commun, 2020. 11(1): p. 4113.

4. Damgaci, S., et al., Hypoxia and acidosis: immune suppressors and therapeutic targets. Immunology, 2018. 154(3): p. 354–362.

5. Ibrahim-Hashim, A. and V. Estrella, Acidosis and cancer: from mechanism to neutralization. Cancer Metastasis Rev, 2019. 38(1-2): p. 149–155.

6. Lardner, A., The effects of extracellular pH on immune function. J Leukoc Biol, 2001. 69(4): p. 522–30.

7. Bohn, T., et al., Tumor immunoevasion via acidosis-dependent induction of regulatory tumor-associated macrophages. Nat Immunol, 2018. 19(12): p. 1319–1329.

8. Davern, M., et al., Acidosis significantly alters immune checkpoint expression profiles of T cells from oesophageal adenocarcinoma patients. Cancer Immunol Immunother, 2023. 72(1): p. 55–71.

9. Potzl, J., et al., Reversal of tumor acidosis by systemic buffering reactivates NK cells to express IFN-gamma and induces NK cell-dependent lymphoma control without other immunotherapies. Int J Cancer, 2017. 140(9): p. 2125–2133.

10. Silva, A.S., et al., The potential role of systemic buffers in reducing intratumoral extracellular pH and acid-mediated invasion. Cancer Res, 2009. 69(6): p. 2677–84.

11. Robey, I.F., et al., Bicarbonate increases tumor pH and inhibits spontaneous metastases. Cancer Res, 2009. 69(6): p. 2260–8.

12. Persi, E., et al., Systems analysis of intracellular pH vulnerabilities for cancer therapy. Nat Commun, 2018. 9(1): p. 2997.

13. Bellone, M., et al., The acidity of the tumor microenvironment is a mechanism of immune escape that can be overcome by proton pump inhibitors. Oncoimmunology, 2013. 2(1): p. e22058.

14. Calcinotto, A., et al., Modulation of microenvironment acidity reverses anergy in human and murine tumor-infiltrating T lymphocytes. Cancer Res, 2012. 72(11): p. 2746–56.

15. Ferrari, S., et al., Proton pump inhibitor chemosensitization in human osteosarcoma: from the bench to the patients’ bed. J Transl Med, 2013. 11: p. 268.

16. Wang, B.Y., et al., Intermittent high dose proton pump inhibitor enhances the antitumor effects of chemotherapy in metastatic breast cancer. J Exp Clin Cancer Res, 2015. 34(1): p. 85.

17. Pilon-Thomas, S., et al., Neutralization of Tumor Acidity Improves Antitumor Responses to Immunotherapy. Cancer Res, 2016. 76(6): p. 1381–90.

18. Gillies, R.J., et al., Back to basic: Trials and tribulations of alkalizing agents in cancer. Front Oncol, 2022. 12: p. 981718.

19. Beauchemin, N. and A. Arabzadeh, Carcinoembryonic antigen-related cell adhesion molecules (CEACAMs) in cancer progression and metastasis. Cancer Metastasis Rev, 2013. 32(3-4): p. 643–71.

20. Burgos, M., et al., Prognostic value of the immune target CEACAM6 in cancer: a meta-analysis. Ther Adv Med Oncol, 2022. 14: p. 17588359211072621.

21. Tian, B., et al., Production and characterization of a camelid single domain antibody-urease enzyme conjugate for the treatment of cancer. Bioconjug Chem, 2015. 26(6): p. 1144–55.

22. Piha-Paul, S., et al., A Phase 1, Open-Label, Dose-Escalation Study of L-DOS47 in Combination With Pemetrexed Plus Carboplatin in Patients With Stage IV Recurrent or Metastatic Nonsquamous NSCLC. JTO Clin Res Rep, 2022. 3(11): p. 100408.

23. An, F., et al., Carcinoembryonic Antigen Related Cell Adhesion Molecule 6 Promotes Carcinogenesis of Gastric Cancer and Anti-CEACAM6 Fluorescent Probe Can Diagnose the Precancerous Lesions. Front Oncol, 2021. 11: p. 643669.

24. Kurlinkus, B., et al., CEACAM6’s Role as a Chemoresistance and Prognostic Biomarker for Pancreatic Cancer: A Comparison of CEACAM6’s Diagnostic and Prognostic Capabilities with Those of CA19-9 and CEA. Life (Basel), 2021. 11(6).

25. Bian, J. and K. Almhanna, Pancreatic cancer and immune checkpoint inhibitors-still a long way to go. Transl Gastroenterol Hepatol, 2021. 6: p. 6.

26. Marabelle, A., et al., Efficacy of Pembrolizumab in Patients With Noncolorectal High Microsatellite Instability/Mismatch Repair-Deficient Cancer: Results From the Phase II KEYNOTE-158 Study. J Clin Oncol, 2020. 38(1): p. 1–10.

27. O’Reilly, E.M., et al., Durvalumab With or Without Tremelimumab for Patients With Metastatic Pancreatic Ductal Adenocarcinoma: A Phase 2 Randomized Clinical Trial. JAMA Oncol, 2019. 5(10): p. 1431–1438.

28. Balachandran, V.P., G.L. Beatty, and S.K. Dougan, Broadening the Impact of Immunotherapy to Pancreatic Cancer: Challenges and Opportunities. Gastroenterology, 2019. 156(7): p. 2056–2072.

29. Bogdanov, A., et al., Tumor acidity: From hallmark of cancer to target of treatment. Front Oncol, 2022. 12: p. 979154.

30. Ettinger, S.N., et al., Urea as a recovery marker for quantitative assessment of tumor interstitial solutes with microdialysis. Cancer Res, 2001. 61(21): p. 7964–70.

31. Kabacaoglu, D., et al., Immune Checkpoint Inhibition for Pancreatic Ductal Adenocarcinoma: Current Limitations and Future Options. Front Immunol, 2018. 9: p. 1878.

32. Feng, M., et al., PD-1/PD-L1 and immunotherapy for pancreatic cancer. Cancer Letters, 2017. 407: p. 57–65.

33. Mucileanu, A., R. Chira, and P.A. Mircea, PD-1/PD-L1 expression in pancreatic cancer and its implication in novel therapies. Med Pharm Rep, 2021. 94(4): p. 402–410.

34. Pillai, S.R., et al., Causes, consequences, and therapy of tumors acidosis. Cancer Metastasis Rev, 2019. 38(1-2): p. 205–222.

35. Ibrahim-Hashim, A., et al., Systemic buffers inhibit carcinogenesis in TRAMP mice. J Urol, 2012. 188(2): p. 624–31.

36. Estrella, V., et al., Acidity generated by the tumor microenvironment drives local invasion. Cancer Res, 2013. 73(5): p. 1524–35.

37. Ibrahim Hashim, A., et al., Reduction of metastasis using a non-volatile buffer. Clin Exp Metastasis, 2011. 28(8): p. 841–9.

38. Ibrahim-Hashim, A., et al., Tris-base buffer: a promising new inhibitor for cancer progression and metastasis. Cancer Med, 2017. 6(7): p. 1720–1729.

39. Pilot, C., A. Mahipal, and R. Gillies, Buffer Therapy → Buffer Diet. Journal of Nutrition & Food Sciences, 2018. 08.

40. Hamaguchi, R., R. Narui, and H. Wada, Effects of Alkalization Therapy on Chemotherapy Outcomes in Metastatic or Recurrent Pancreatic Cancer. Anticancer Res, 2020. 40(2): p. 873–880.

41. Hamaguchi, R., et al. Improved Chemotherapy Outcomes of Patients With Small-cell Lung Cancer Treated With Combined Alkalization Therapy and Intravenous Vitamin C. Cancer diagnosis & prognosis, 2021. 1, 157–163 DOI: 10.21873/cdp.10021.

42. Hamaguchi, R., et al., Clinical review of alkalization therapy in cancer treatment. Front Oncol, 2022. 12: p. 1003588.

43. Ramlau, R., et al., P2.06-006 Phase I/II Dose Escalation Study of L-DOS47 as a Monotherapy in Non-Squamous Non-Small Cell Lung Cancer Patients: Topic: Phase I/II Trials. Journal of Thoracic Oncology, 2017. 12(1, Supplement): p. S1071–S1072.

44. Ibrahim-Hashim, A., et al., Defining Cancer Subpopulations by Adaptive Strategies Rather Than Molecular Properties Provides Novel Insights into Intratumoral Evolution. Cancer Res, 2017. 77(9): p. 2242–2254.

45. Jardim-Perassi, B.V., et al., Intraperitoneal Delivery of Iopamidol to Assess Extracellular pH of Orthotopic Pancreatic Tumor Model by CEST-MRI. Contrast Media Mol Imaging, 2023. 2023: p. 1944970.

46. Bates, D., et al., Fitting Linear Mixed-Effects Models Using lme4. Journal of Statistical Software, 2015. 67(1): p. 1 –48.

47. Kuznetsova, A., P.B. Brockhoff, and R.H.B. Christensen, lmerTest Package: Tests in Linear Mixed Effects Models. Journal of Statistical Software, 2017. 82(13): p. 1–26.

48. Team, R.C., R: A language and environment for statistical computing. 2020, R Foundation for Statistical Computing: Vienna, Austria.

49. Searle, S.R., F.M. Speed, and G.A. Milliken, Population Marginal Means in the Linear Model: An Alternative to Least Squares Means. The American Statistician, 1980. 34(4): p. 216–221.

