## Supplemental for "L-DOS47 enhances response to immunotherapy in pancreatic cancer tumor"

### Supplemental Figures

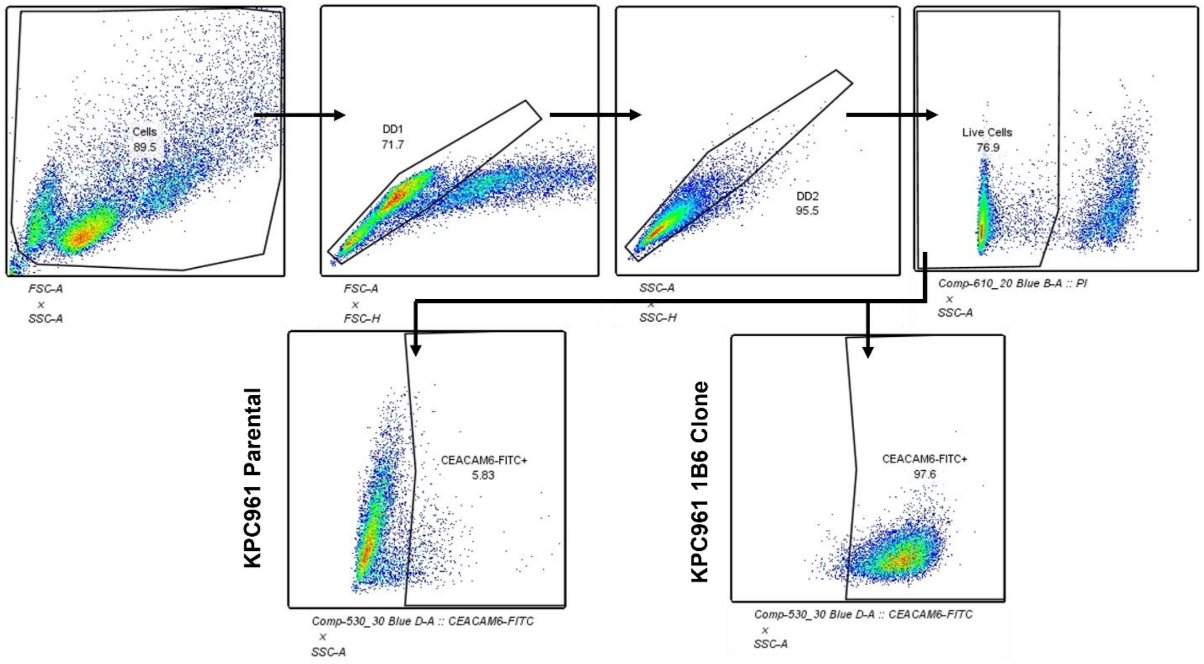


**Supplemental Fig 1.** Gating strategy used for flow cytometer analysis. FACS plots are shown as representative example of the gating strategy used to quantify the CEACAM6 expression in KPC961 clone 1B6 cells after transduction of human CEACAM6.

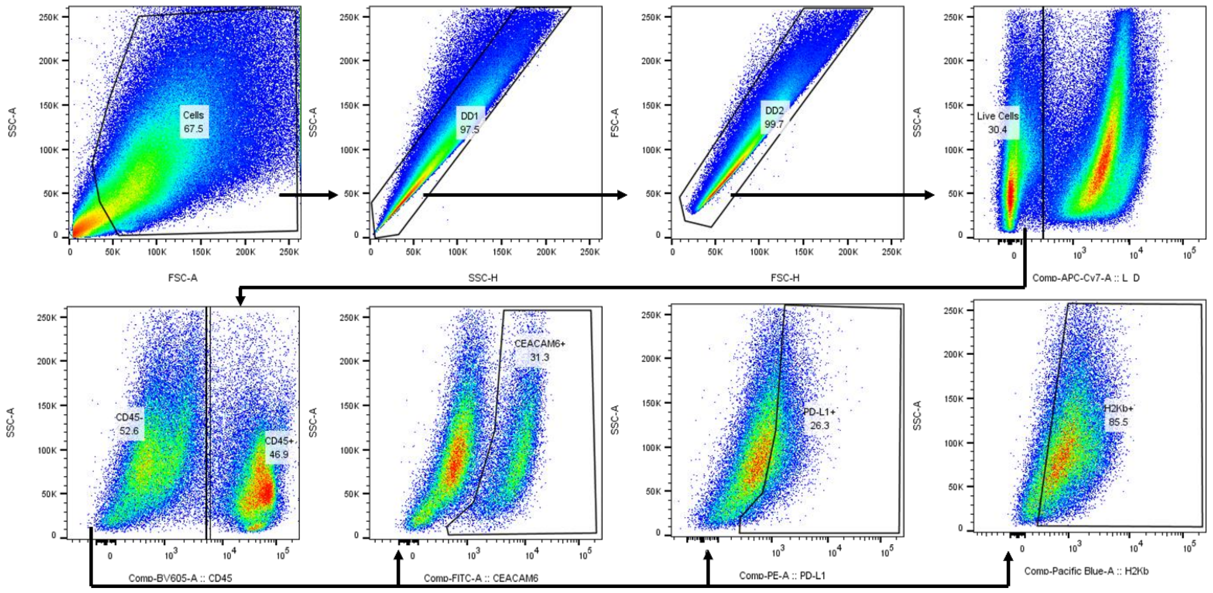


**Supplemental Fig 2. Gating strategy used for flow cytometer analysis.** FACS plots are shown as representative example of the gating strategy used to quantify the expression of CEACAM6, PDL-1, and H2Kb in inoculated KPC961-1B6 tumors

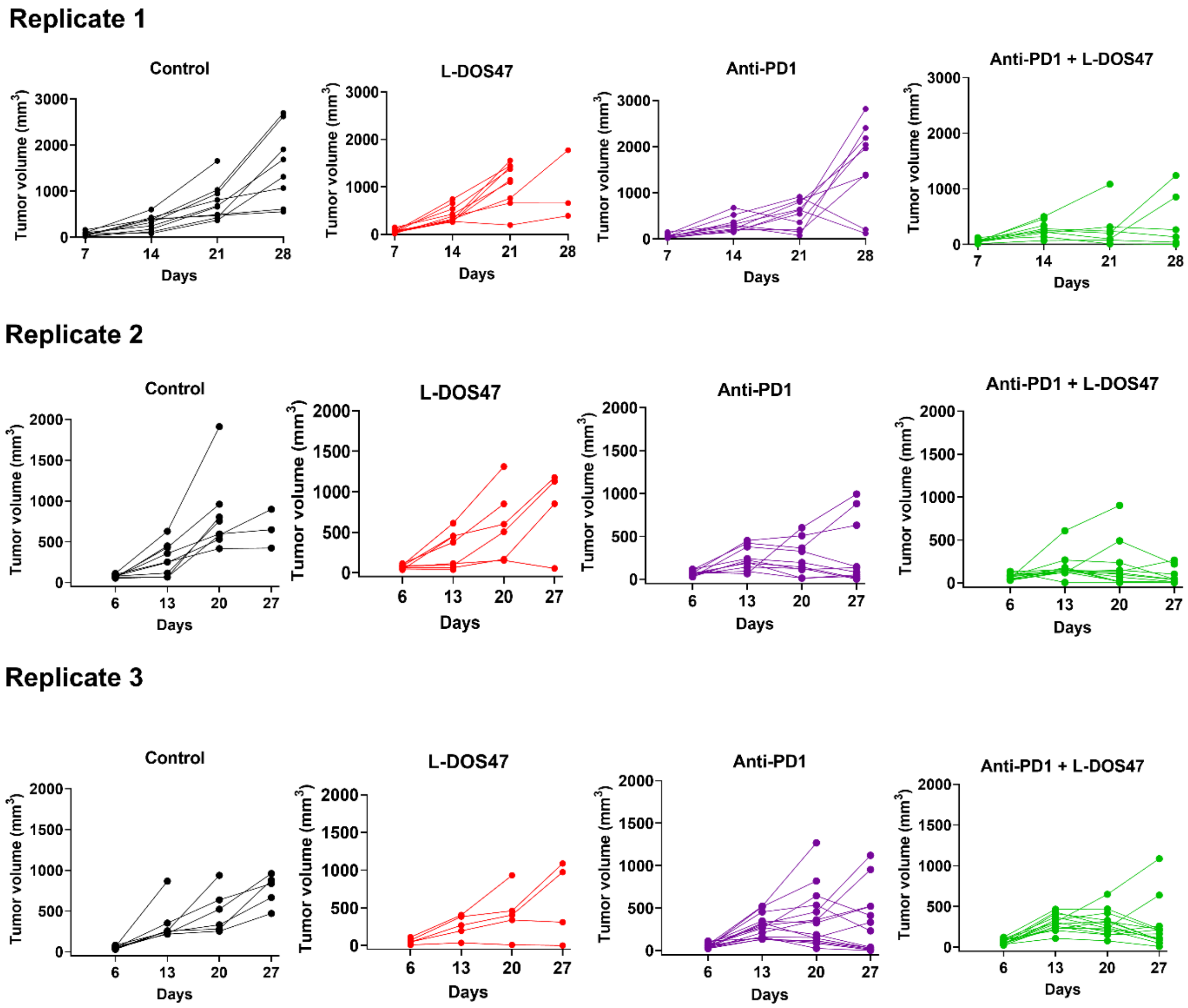


**Supplemental Fig 3. Individual tumor growth for each therapy group for replicates 1, 2 and 3.** Number of mice per group in each replicates were: Replicate 1 Control=10, anti-PD1=10, L-DOS47 = 10, anti-PD1 + L-DOS47 =9; Replicate 2 Control = 9, anti-PD1=10, L-DOS47 =8, anti-PD1 + L-DOS47 =12; and replicate 3 Control =7, anti-PD1 =15, L-DOS47 =5, anti-PD1 + L-DOS47 =14.

**Table S1.** Coefficients from the linear mixed effects model for tumor growth by treatment arm for Experiments 1, 2 and 3 combined. The reference treatment arm (represented by the intercept) is Anti-PD1 + L-DOS47.

| Fixed effects | Estimate | Std. Error | df | t value | Pr(>\|t\|) |
|---|---|---|---|---|---|
| (Intercept) | 6.43 | 0.30 | 474.58 | 21.06 | < 2e-16 |
| Anti-PD1 | -0.36 | 0.42 | 469.39 | -0.86 | 0.38 |
| L-DOS47 | -0.88 | 0.47 | 462.44 | -1.85 | 0.06 |
| Control | -1.12 | 0.46 | 468.34 | -2.42 | 0.01 |
| day | 0.02 | 0.01 | 413.99 | 1.51 | 0.13 |
| Anti-PD1: day | 0.04 | 0.02 | 414.16 | 2.17 | 0.02 |
| LDOS47: day | 0.11 | 0.02 | 414.01 | 4.77 | 2.50e-06 |
| Control: day | 0.14 | 0.02 | 414.16 | 5.96 | 5.18e-09 |
